# High resolution atlas of the venous brain vasculature from 7 T quantitative susceptibility maps

**DOI:** 10.1101/444349

**Authors:** Julia Huck, Yvonne Wanner, Audrey P. Fan, Anna-Thekla Jäger, Sophia Grahl, Uta Schneider, Arno Villringer, Christopher J. Steele, Christine L. Tardif, Pierre-Louis Bazin, Claudine J. Gauthier

**Affiliations:** Concordia University, Department of Physics, Montreal, Canada; Universität Stuttgart, Stuttgart, Germany; Stanford University, Stanford, United States; Max-Planck-Institut fur Kognitions- und Neurowissenschaften, Leipzig, Germany; Clinic for Cognitive Neurology, University of Leipzig, Leipzig, Germany; Leipzig University Medical Centre, IFB Adiposity Diseases, Leipzig, Germany; Leipzig University Medical Centre, Collaborative Research Centre 1052-A5, Leipzig, Germany; Concordia University, Department of Psychology, Montreal, Canada; McGill University, Department of Biomedical Engineering, Montreal, Canada; Montreal Neurological Institute, Montreal, Canada; University of Amsterdam, Faculty of Social and Behavioural Sciences, Amsterdam, Netherlands; Montreal Heart Institute, Montreal, Canada

**Keywords:** venous vasculature, QSM, UHF-MRI, vein atlas, vein segmentation, cerebral vasculature

## Abstract

The vascular organization of the human brain can determine neurological and neurophysiological functions, yet thus far it has not been comprehensively mapped. Aging and diseases such as dementia are known to be associated with changes to the vasculature and normative data could help detect these vascular changes in neuroimaging studies. Furthermore, given the well-known impact of venous vessels on the blood oxygen level dependent (BOLD) signal, information about the common location of veins could help detect biases in existing datasets. In this work, a quantitative atlas of the venous vasculature using quantitative susceptibility maps (QSM) acquired with a 0.6 mm isotropic resolution is presented. The Venous Neuroanatomy (VENAT) atlas was created from 5 repeated 7 Tesla MRI measurements in young and healthy volunteers (n = 20, 10 females, mean age = 25.1 ± 2.5 years) using a two-step registration method on 3D segmentations of the venous vasculature. This cerebral vein atlas includes the average vessel location, diameter (mean: 0.84 ± 0.33 mm) and curvature (0.11 ± 0.05 mm^−1^) from all participants and provides an in vivo measure of the angio-architectonic organization of the human brain and its variability. This atlas can be used as a basis to understand changes in the vasculature during aging and neurodegeneration, as well as vascular and physiological effects in neuroimaging.

## Introduction

Cerebral arteries are known to be affected in aging and diseases such as atherosclerosis and dementia (Peters 2006; Brown and Thore 2011). In part because of this, arteries have long been the main focus of research on the cerebral vasculature. The venous vasculature is more challenging to characterize due to greater variability between subjects (Browning 1884; Duvernoy et al. 1981; Bernier et al. 2018), but there is an increasing recognition that valuable information about cerebral physiology and brain function in health and disease could be gleaned from a more thorough understanding of veins.

Veins have received increased attention over the past decades as they are a significant source of both contrast and bias in gradient echo (GE) blood oxygenation level dependent (BOLD) functional magnetic resonance imaging (fMRI) (Seiyama et al. 2004; Gagnon et al. 2015), especially at low field strengths (Donahue et al. 2011). But while different aspects of venous physiology have been explored in the context of the BOLD signal, venous structure has only received a limited amount of attention (Turner 2002; Gagnon et al. 2015). One of the issues of concern is that the GE BOLD contrast is known to be sensitive to draining veins, which introduce a bias in the measured location of brain activation (Turner 2002). This is because the BOLD signal is based on local changes in deoxyhemoglobin (dHb) concentration from dilution with oxyhemoglobin during the hemodynamic response that accompanies changes in neural activity (Logothetis and Wandell 2004). While the BOLD signal is thought to arise partly from the parenchymal response, it has been shown to also have a significant intravascular component, especially at the lower field strengths of 1.5 or 3 Tesla (T) used in most functional MRI studies (Boxerman et al. 1995; Donahue et al. 2011). This is problematic since it is estimated that a vein with a diameter of 0.6 mm is able to drain 125 mm² of cortex (Turner 2002). This influence of draining veins generates uncertainty in the exact location of brain activity as measured by the BOLD response, but also biases the amplitude of the response measured as draining veins contain more dHb and may therefore contribute the voxels with the highest signal change amplitude. These venous voxels may then be detected as the most significant areas of signal change instead of tissue (Boubela et al. 2015). There is currently no established method to identify these voxels with large veins in vivo. However, a venous atlas could in the future be used as a prior to identify the voxels most likely to have a large venous contribution in functional MRI studies, and would allow the measurement of orientation-related biases.

It is also increasingly recognized that factors such as aging, injury, diseases such as stroke and dementia, lifestyle changes and physiological training can have a profound impact on brain vascular structure and oxidative metabolism (Kramer et al. 2006; Pathak et al. 2011; Voss et al. 2011). Structural changes in the cerebral vasculature have been shown to occur in post-mortem studies of aging, Alzheimer’s disease (AD), cerebral venous thrombosis (CVT) (Towbin 1973; Villringer et al. 1989; Einhäupl et al. 1991; Vogl et al. 1994) and leukoaraiosis (Brown and Thore 2011). A reduced vascular density and increased tortuosity has been observed in aging, and these changes are thought to underlie the development of the ubiquitous white matter lesions (Peters 2006). While these changes have predominantly been studied in terms of the arterial vasculature, Brown et al. report an increased wall thickness in veins and venules of the periventricular white matter with normal aging and a greater degree of wall thickening in patients with leukoaraiosis (Brown and Thore 2011). Furthermore, an increased tortuosity has been associated with leukoaraiosis (Wahlund et al. 2001) and has been shown to be detectable using susceptibility-weighted imaging (SWI) at 7 T (Shaaban et al. 2017). The small size of the veins affected by many neurological disorders poses a challenge for in vivo imaging however, especially at the field strengths typically used for MRI studies of aging and disease. An atlas of the venous vasculature could help solve some of these issues by providing a strong prior for venous segmentation of susceptibility images and normative information against which to compare aging and disease-related changes.

Magnetic susceptibility is an intrinsic property of tissues that describes the response of its atoms in a large magnetic field, such as the paramagnetic effect of iron. This property can be imaged using the Quantitative Susceptibility Mapping (QSM) technique, which has been shown to be primarily sensitive to myelin, iron deposition and dHb in the brain (Langkammer et al. 2012; Stüber et al. 2014; Fan et al. 2014; Wang and Liu 2015). QSM can be used to image veins, since the presence of paramagnetic dHb molecules creates a difference in magnetic susceptibility in venous blood relative to surrounding tissue. Furthermore, the quantitative susceptibility in each venous voxel is directly related to the Oxygen Extraction Fraction (OEF) and the Cerebral Metabolic Rate of Oxygen (CMRO_2_) through Fick’s principle (Fan et al. 2014; Serres et al. 2015). Therefore, QSM is a powerful technique for studying the venous network as it not only provides information about venous structure, but also about local metabolism.

Recognition of the importance of the brain’s venous network combined with recent developments in susceptibility imaging has already led to the creation of two probabilistic venous atlases (Ward et al. 2018; Bernier et al. 2018). QSM can however provide additional information about venous properties, which we seek to leverage here. In this paper we introduce the Venous Neuroanatomy (VENAT) atlas. This atlas provides several advantages over existing venous atlases, since it was acquired with a higher field strength (higher SNR), has a higher image resolution (0.6 mm isotropic) and uses multiple repetitions to increase SNR. This image resolution allows detection of vessels up to 0.3 mm in diameter. The VENAT atlas is open source and available on figshare (https://figshare.com/articles/VENAT_Probability_map_nii_gz/7205960). This atlas was built using an optimized approach through registration of vascular segmentations and using five repeated QSM images per participant. The VENAT atlas includes a partial volume (PV) map, which shows vessel location, maps of the diameter, curvature of vessels, and venous density maps.

## Methods

### Participants

MRI acquisitions were performed in twenty young, right-handed participants between the ages of 22 and 30 years (average ± standard deviation (SD) of 25.05 ± 2.48 years, 10 females). None of the participants had a history of neurological disorders or currently suffered from psychiatric disorders as indicated by self-report and a structured clinical interview. The study was approved by the Ethics Committee of the University of Leipzig and was conducted in accordance with the Declaration of Helsinki. All participants gave written informed consent prior to the beginning of the study.

### Scan Parameters

Acquisitions were completed on a 7 T Siemens Magnetom MR system (Siemens Healthcare, Erlangen, Germany) with a 32 channel Nova head coil (NOVA Medical Inc., Wilmington MA, USA). Two sequences were acquired to generate the vascular atlas: a multi-echo 3D GE FLASH (Haase et al. 1986) with two echo times (TE) and a whole brain T1 map using the MP2RAGE technique (Marques et al. 2010). Auto-align was used at the beginning of each session to ensure a comparable placement of the acquired image between time points. Additionally, dielectric pads were placed at the sides of their head to enhance signal in the temporal lobes and the cerebellum (O’Reilly et al. 2016).

The multi-echo 3D GE FLASH image was acquired with a 0.6 mm isotropic resolution; flow compensation along all three axes for the first echo; repetition time (TR) = 29 ms; TE1 / TE2 = 8.16 / 18.35 ms; matrix = 260×320×256; GRAPPA acceleration = 3; bandwidth = 250 Hz/Px; and time of acquisition (TA)=14:22 min. Phase and magnitude information was saved separately for each receive channel. The whole brain MP2RAGE was acquired with a 0.7 mm isotropic resolution and the following parameters: TR = 5000 ms; TE = 2.45 ms; matrix= 320×320×240; inflow time 1/2 = 900 / 2750 ms; Flip angle 1 / 2 = 5° / 3°; bandwidth = 250 Hz/Px; and TA = 10:57 min. The resulting estimated quantitative T1 map was used for registration. Additionally, low resolution GE phase reference images were acquired to estimate phase offsets between receive channels for optimal coil combination of the high resolution phase images (Hammond et al. 2008; Deistung et al. 2013). The scan parameters were TR = 18 ms, TE = 4.08 / 9.18 ms, matrix = 128×128×80, flip angle = 10°, 2 mm isotropic resolution, bandwidth = 300 Hz/Px, and TA = 3:24 min.

Identical acquisitions were performed five times over a period of three weeks on each participant as part of an interventional motor learning study, with 20 min of training for 5 days. Since we did not expect changes in macrovascular venous structure as a consequence of this learning paradigm, all five acquisitions were used for generating the atlas.

### Registration to standard stereotactic MNI-space

Processing was done using Medical Image Processing, Analysis and Visualization (MIPAV) 7.4.0 [http://mipav.cit.nih.gov/] and the CBS High-Res Brain Processing Tools for MIPAV (CBSTools) [https://www.nitrc.org/projects/cbs-tools/], except when specified. At each time point, a dedicated skull stripping algorithm was applied to the MP2RAGE images (Bazin et al. 2016). The skull stripped images were then registered rigidly to the MNI152 0.5 mm template to avoid deforming individual anatomy. The second echo image of the 3D GE sequence, which provided good anatomical contrast, was registered linearly to the quantitative T1 MP2RAGE image from the same time point. Linear registrations were done using FSL FLIRT (Fischer and Modersitzki 2003) with normalized mutual information and 6 Degrees of Freedom (DoF). The deformation field from the registration from the MP2RAGE images to MNI space and from 3D GE to MP2RAGE were concatenated and applied to the first echo from the 3D GE sequence to transform these images from native space into MNI space.

### Quantitative Susceptibility Map reconstruction

The phase images from each channel were recombined offline using the phase estimates from the low-resolution field maps to ensure a high phase image quality for QSM reconstruction. The combined magnitude (**Fig. *1*b**) and phase (**Fig. *1*a**) of the first, flow-compensated echo (TE = 8.16ms) were used as input for QSM reconstruction. A brain mask was generated by building an average of the MP2RAGE images from all five time points for each participant in MNI space. This average brain mask was generated to ensure a consistent brain size over all time points. The average brain mask was transformed into individual space for each participant and time point by applying the inverse transformation field from the registration of the 3D GE images to MNI space.

**Fig. 1:**
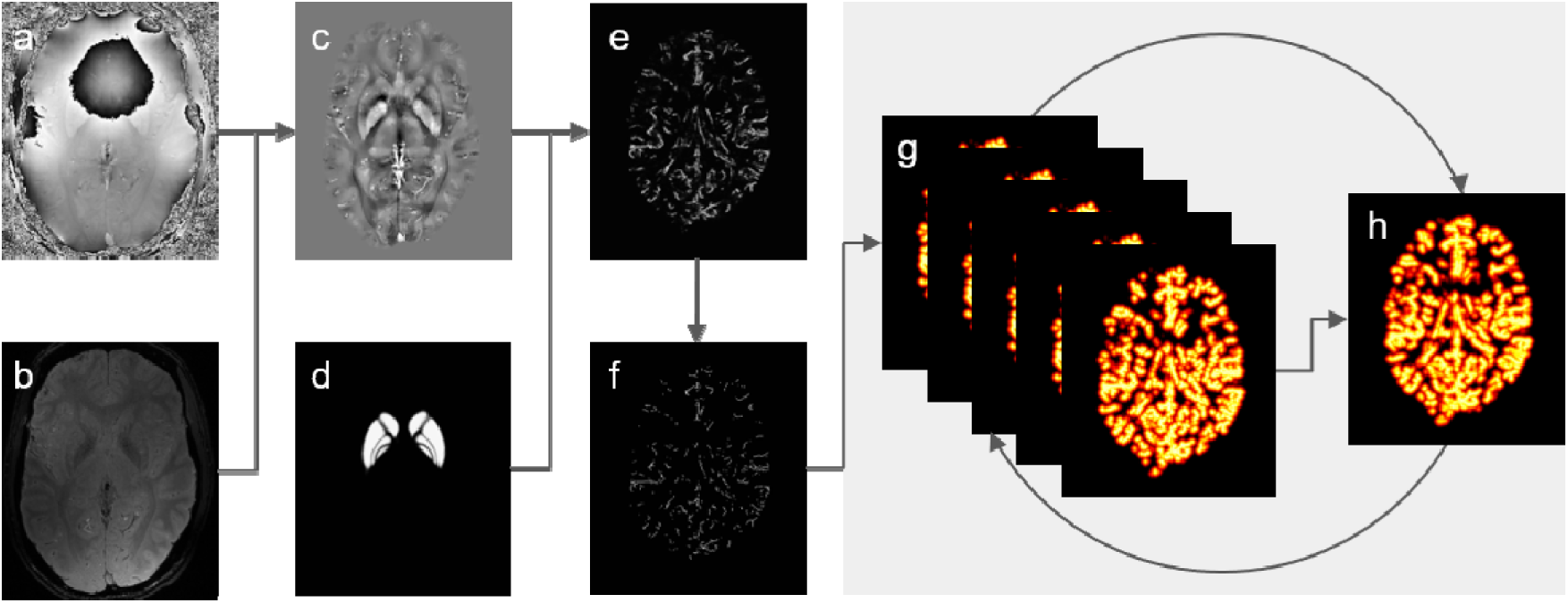
Pipeline for the single participant average; **a)** Combined phase (TE = 8.16 ms); **b)** Combined magnitude (TE = 8.16 ms); **c)** QSM; **d)** ATAG atlas; **e)** probability map of the vessel segmentation; **f)** skeletonized vessels; **g)** probability distance map (up to 15 mm) of all time points; **h)** average of probability distance maps.

The brain mask was eroded by 8 voxels to avoid artifacts from the large susceptibility difference at air-tissue interfaces during QSM reconstruction while preserving as much brain volume as possible. QSM maps were created using the Fast L1-Regularized QSM with Magnitude Weighting and SHARP background filtering toolbox (Bilgic et al. 2014) (**Fig. *1*c**) in MATLAB R2016b (MathWorks, Inc., Natick, Massachusetts, United States. This QSM reconstruction method was chosen to ensure an optimal parameter estimation for each participant individually using the L-curve method. Magnitude weighting was used to enhance contrast around veins. The standard toolbox parameters were only modified to change the field strength parameter from 3 T to 7 T.

### Vessel Segmentation

Vessels were segmented using a multiscale vessel filter (Bazin et al. 2016) on the QSM image from each time point. This filter estimates image edges recursively and uses a global probability diffusion scheme to link nearby vascular voxels into vessel branches. This vessel filter generates a probability map, a partial volume map and a diameter map as output. It estimates vessel diameter based on PV of the smaller vessels in a multiscale fashion. This diameter estimation technique can detect diameters half the voxel size of the resolution using the approximation described by Woerz et al. (Woerz and Rohr 2004). These diameter maps depict a one voxel thick centerline with diameter coded as the intensity value. This filter has been found to be particularly sensitive to small vessels, see Bazin et al., (2016) for an evaluation. However, due to the high iron content of the smaller basal ganglia nuclei, the vessel filter also detects small basal ganglia nuclei as large vessels. To prevent this erroneous segmentation, the Atlas of The bAsal Ganglia (ATAG) atlas of 30 young participants, was used as a negative prior to modify the filter (**Fig. *1*d**) (Keuken et al. 2014). The negative prior included the non-linear right and left masks of the striatum, external segment of the globus pallidus, internal segment of the globus pallidus, red nucleus, subthalamic nucleus, substantia nigra, and periaqueductal grey, combined into one mask in MNI space. To transform the ATAG mask from MNI space into individual space, the inverse deformation fields from the 3D GE to MNI space registration were applied. Vessel segmentation with the multiscale filter was done in native space to avoid segmentation errors due to registration of the QSM images into MNI space. In order to calculate an average of the diameter images even when the alignment across time points and participants is imperfect, the diameter maps were inflated by 2 mm to create a 2 mm skeleton of the vasculature. The output images of the vessel filter (vessel probability map, PV map and the inflated diameter) were then registered into the MNI152 space by applying the transformation matrices of the 3D GE images into the MNI space.

### Atlas Construction

#### Single Participant Average

The atlas was created by registering the segmented vessels from all time points for each participant to each other and calculating the mean of these images. This type of registration is challenging given the small size of cortical vessels and variability in vessel location across participants or due to registration errors across participants. To prevent this loss of information, the segmented vessels in MNI space (**Fig. *1*e**) were reduced to their centerline by applying a skeletonization algorithm adapted from Bouix et al. (Bouix et al. 2005) on the probability maps, with a boundary and skeleton threshold of 0.5 (**Fig. *1*f**). To increase the probability of an overlap of the vessel location, a distance map up to 15 mm (unless the veins were located closer to each other) of these centerlines was calculated. These maps were then converted to probability distance maps (**Fig. *1*g**) (i.e. a probability of 1 at the centerline with decreasing values with increasing distance). This allowed the construction of an average vessel location despite some variability in location. Curvature was calculated from the skeletons of the probability maps in MNI space. The algorithm for calculating the curvature was adapted from An et al. (An et al. 2011). An et al. computes the curvature based on discrete curves. The curvature is geometrically defined as the inverse radius of the circle at every point of the curve. The curvature maps are depicted as a skeleton with their corresponding curvature as intensity value. Similar to the diameter maps, these curvature maps were also inflated by 2 mm to ensure an overlap during the averaging process even if the alignment is imperfect.

The probability distance maps in MNI space of each time point were registered to each other using the SyN module of Advanced Normalization Tools (ANTS) (Avants et al. 2009). The reference image used for this registration was the average of the participant’s probability distance maps in MNI space (**Fig. *1*h**). The SyN algorithm was run with the standard parameters as specified in the CBSTools interface with 40 × 50 × 40 iterations at coarse, medium and fine scales, a three voxel gaussian blurring kernel, and SyN parameter set to 0.25. Registration was done iteratively with each step taking as input the probability distance maps from the previous registration with their resulting average as a reference (**Fig. *1*g-h**). After each registration step the transformation matrices of the probability distance maps were also applied on the PV maps, the inflated diameters and inflated curvature maps and their corresponding averages were calculated. To follow the improvement from each registration step, the standard deviation of each registered PV image to the average PV map was calculated. Four registration steps were necessary to bring the average improvement for each participant across time points below 10%.

To display the range of our data quality, **Fig. 2** shows the images for the participant with the lowest (denoted as participant low variance (LV)) and highest (high variance (HV)) average standard deviation. For visualization purposes a skeletonization algorithm was then applied on the average registered PV maps with values of 0.33 for the boundary and skeleton threshold to return to a one voxel thick map of the diameter and curvature. The output of the skeletonization was binarized and multiplied with the inflated diameter and curvature average maps of each participant to display diameter and curvature on the skeleton.

**Fig. 2:**
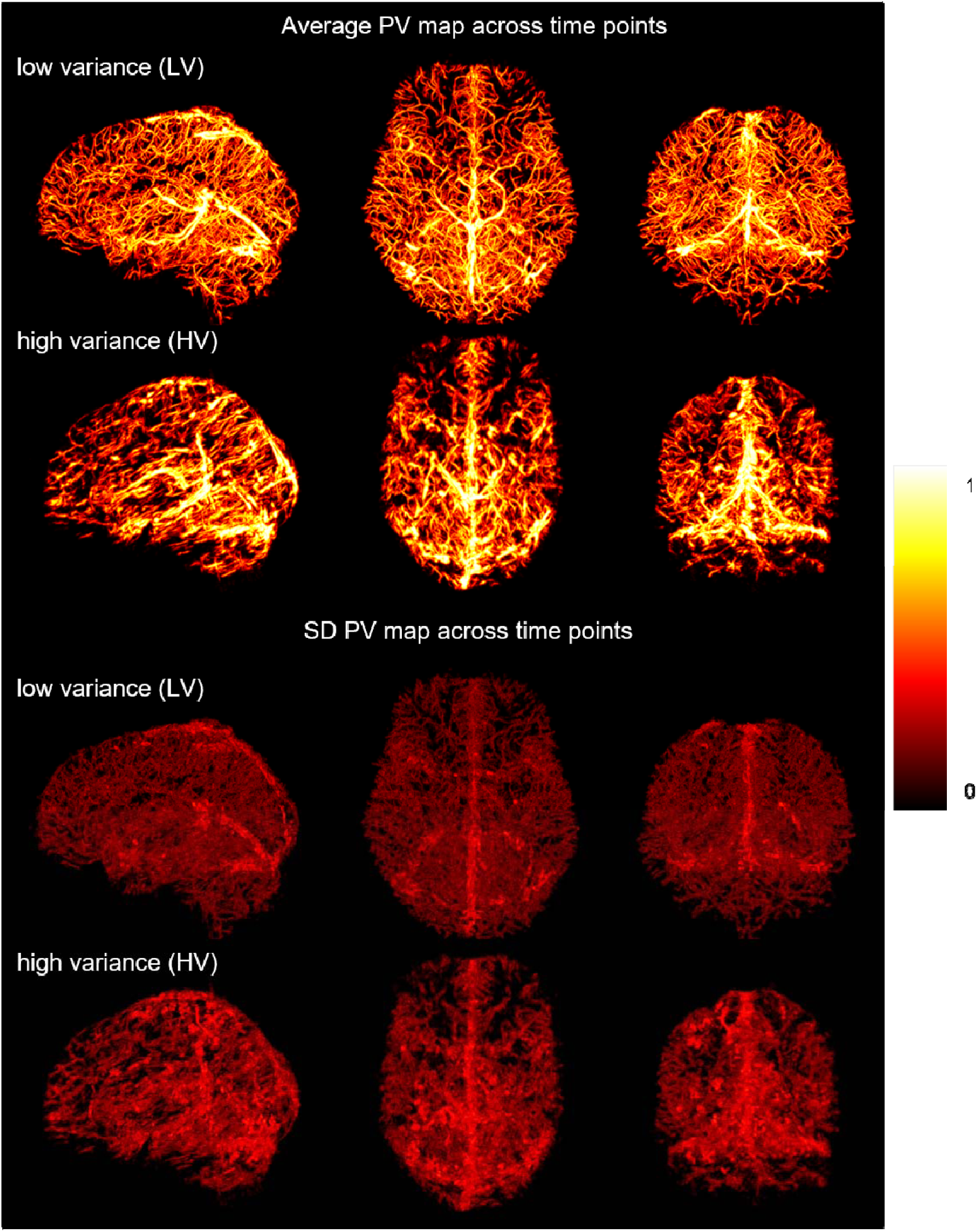
Maximum intensity projection across all slices of the PV maps and the standard deviation; low variance (LV) denotes the participant with the low standard deviation; high variance (HV) denotes the participant with the highest standard deviation. Both participant images are shown in the sagittal, axial and coronal view in MNI152 space.

#### Multi-Participant atlas

The multi-participant atlas was built by combining the individual probability distance averages computed for each individual. These single participant averages (**Fig. 3a**) were combined into a first multi-participant average (**Fig. 3b**). The initial multi-participant average appears blurred as a result of inter-participant variability. To refine the registration, the individual vessel mask that was most similar to the multi-participant average (i.e., with the lowest standard deviation difference) was used as a target and all individual averages non-linearly registered to it with SyN (iteration parameters again set to 50 × 40 × 50). This registration step was done once to ensure a first alignment into the group space, and the resulting registered images were averaged to generate a sharp reference image (**Fig. 3c**) for the following registration steps. After this, multiple iterations using increasingly fine registration parameters (**Fig. 3d-f**) were performed to achieve an accurate alignment of the different participants. Each registration used the average of the previous registration step as a reference image and the registered images of the previous registration step as input to the SyN algorithm of ANTS. As in the single participant average, the transformation matrix was applied after each registration step on the average inflated diameter and curvature maps and the average PV map of the single participants. After each registration an average of these registered maps was calculated.

**Fig. 3:**
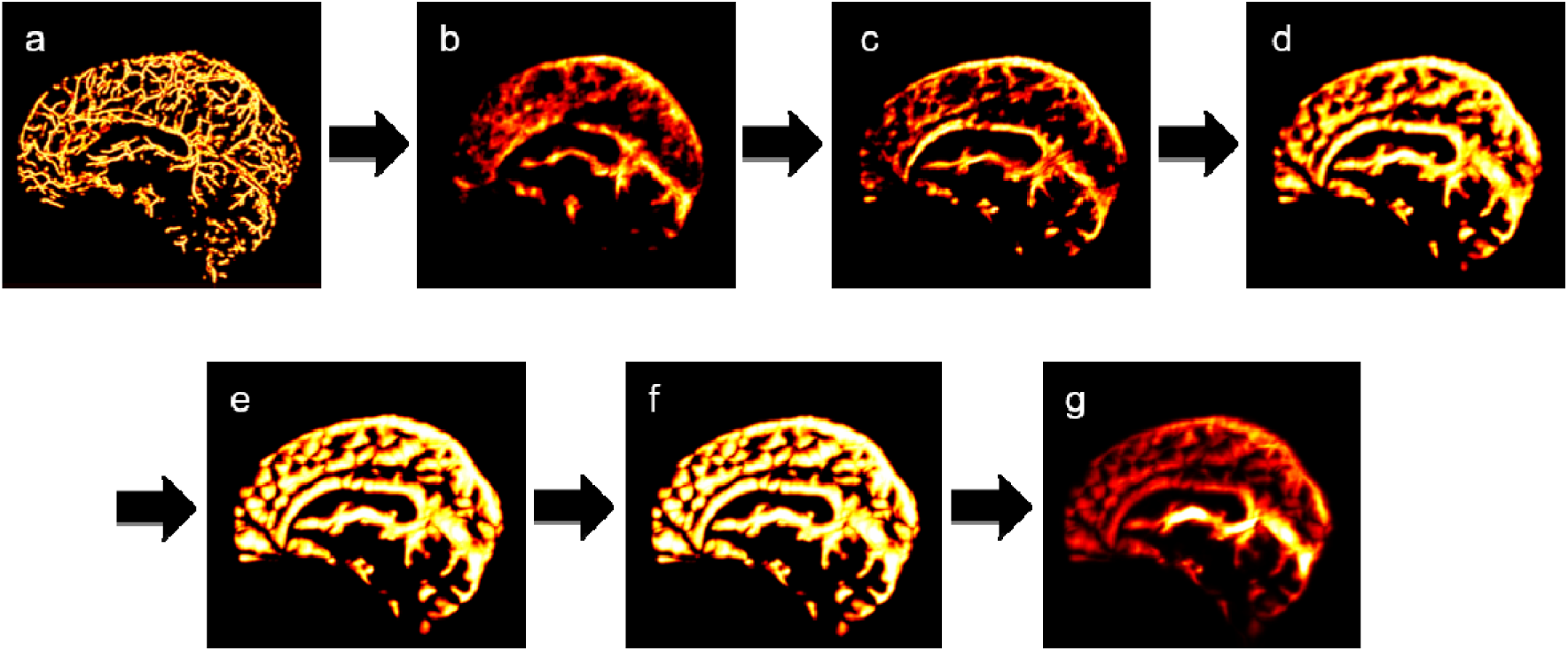
**a)** Venous probability map of the participant with the lowest standard deviation; **b)** Initial multi participant average; **c)** Average after registration to the participant closest to the initial average; **d)** Probability average after 13 coarse registrations; **e)** Probability average after 7 medium registrations; **f)** Probability average after 2 fine registrations; **g)** PV map of the VENAT atlas; all maps are maximum intensity projections across 20 slices starting from the interhemispheric plane.

Each registration step was done until an improvement in standard deviation below 1% was achieved as convergence criterion for each level of registration (**Fig. 4**). Thirteen registration steps were required to meet this criterion at the coarse level (40 × 0 × 0) (**Fig. 3d**), followed by seven medium (40 × 50 × 0) (**Fig. 3e**) and two fine registration steps (40 × 50 × 40) (**Fig. 3f**). After the last registration step, the mean and standard deviation of the PV were calculated. Similar to the single participant averages, a skeletonization algorithm was applied on the final VENAT PV map (**Fig. 3g**) with a boundary/skeleton formation parameter of 0.15. The resulting skeleton was binarized and multiplied with the registered diameter and curvature maps to return to one voxel thick maps for the VENAT atlas.

**Fig. 4:**
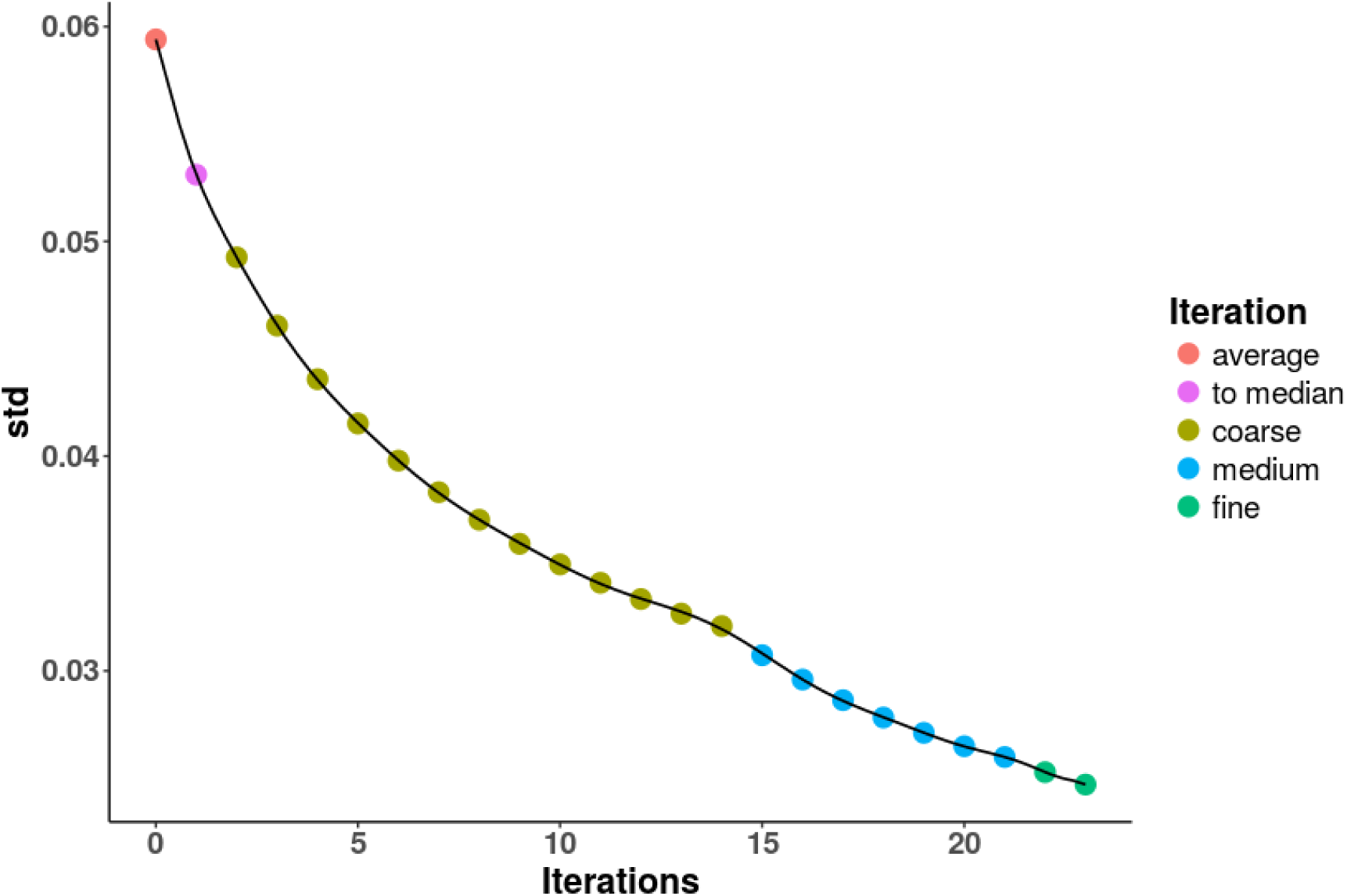
Standard deviation of each iteration step. average denotes the first average of the single participant averages. to median denotes the registration of all participants to the participant closest to the overall average. Iterations were done at each level until the improvement in standard deviation was below 1%.

#### Density Map

A voxel-wise vascular density map was created by using the unthresholded probabilistic vessel maps from the vascular filter. Probabilities larger 0 were binarized and averaged across time points. Only voxels that were detected in at least two time points were kept to reduce the influence of noise. An average across participants of the single participant probability density maps was generated.

Regional density was investigated using both anatomical and functional atlases. Two anatomical atlases were chosen, the MNI and the Harvard-Oxford atlas. The bootstrap analysis of stable clusters (BASC) parcellation (Bellec et al. 2010; Bellec 2013) was used for functional atlas parcellation. The MNI atlas was chosen to estimate densities for each lobe and the Harvard-Oxford and BASC064 atlas for estimating finer grained regional densities based on anatomical and functional parcellations respectively. To maximize sensitivity, we used all venous voxels present in at least two time points in one version of this analysis. In another, we used all venous voxels present in at least three time points to maximize specificity. For each region the mean individual PV was averaged across participants to yield a mean inter-participant average density.

## Results

### Single Participant Average

Examples of inter-session PV averages are shown in **Fig. 2** for the participant with the lowest variance (LV) and highest variance (HV) (mean ± SD of the variance maps: LV: 0.02 ± 0.03; HV: 0.02 ± 0.05) across days. In **Fig. 2**, participant LV demonstrates data quality for our best participant and shows a high density of vessels over the entire brain. HV on the other hand shows a lower density of vessels especially in frontal and temporal areas. Furthermore, LV shows a greater number of smaller cortical veins than HV, which shows mainly larger veins, also appearing blurrier in the image. The standard deviation maps (**Fig. 2**) support these observations as LV shows a lower variance (higher consistency in the vessel diameter) than HV across the brain.

The average diameter across the five time points is given in **Fig. 5** for the same participants as in **Fig. 2**. LV shows more vessels with smaller diameters (mean: 0.67 ± 0.33 mm) compared to HV (mean: 1.01 ± 0.50 mm). The standard deviation (not shown) was on average 0.07 ± 0.07 mm for LV and 0.17 ± 0.14 mm for HV. While most large vessels are not present in the images due to erosion of the mask during QSM reconstruction, part of the sagittal sinus, straight sinus and transverse sinuses are visible in the images. Most visible vessels are smaller cortical vessels however. Participant LV shows a higher vessel density than participant HV, especially in the frontal lobe, and larger numbers of smaller vessels. Furthermore, LV shows a lower variability in vessel diameter, especially for the larger vessels, than HV.

**Fig. 5:**
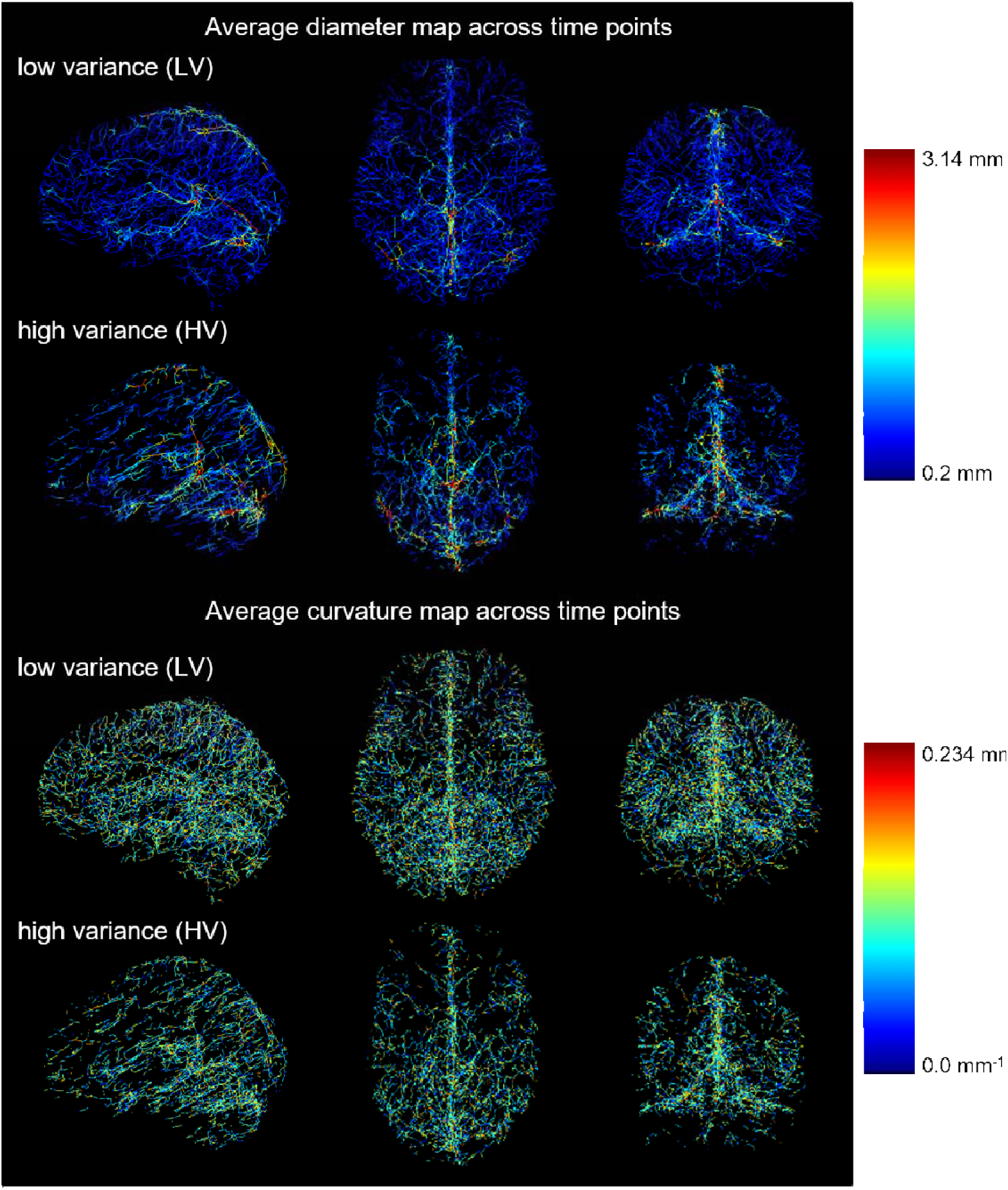
Maximum intensity projection across all slices of the diameter maps and curvature; low variance (LV) denotes the participant with the lowest standard deviation; high variance (HV) denotes the participant with the highest standard deviation. Both participant images are shown in the sagittal, axial and coronal view in MNI152 space.

The average curvature representing the average inverse radius of the circle at every point of the veins is depicted in **Fig. 5**. The figure shows the curvature for the same participants as in **Fig. 2** (LV and HV). Average curvature values were the same for both participants with an average of 0.10 ± 0.04 mm^−1^ (across all voxels identified as veins). To evaluate the consistency across the five time points, the standard deviation was calculated across the brain: respectively 0.06 ± 0.03 mm^−1^ for participant LV and 0.07 ± 0.03 mm^−1^ for participant HV for curvature.

### Multi-participant average

Individual average vessel maps were registered together to create the multi-participant VENAT atlas. **Fig. 6** and **Fig. 7** show the same PV map with a Maximum Intensity Projection (MIP) across 20 slices in the axial view from inferior to superior and in the sagittal view from right to left respectively, to better appreciate the distribution of vessels over the brain. The intensities of the PV images range from 0 to 0.8. Visually, the distribution of vessels was found to be sparser as compared to the single participant averages and to differ across brain regions. **Fig. 8** compares the standard deviation across participants (inter-participant) with the mean standard deviation within participants (intra-participant). The standard deviation ranged from 0 to 0.20 and from 0 to 0.29 for the mean value within participants and across participants, respectively. Standard deviation was found to be smaller within participants than across participants, with larger standard deviations in larger veins.

**Fig. 6:**
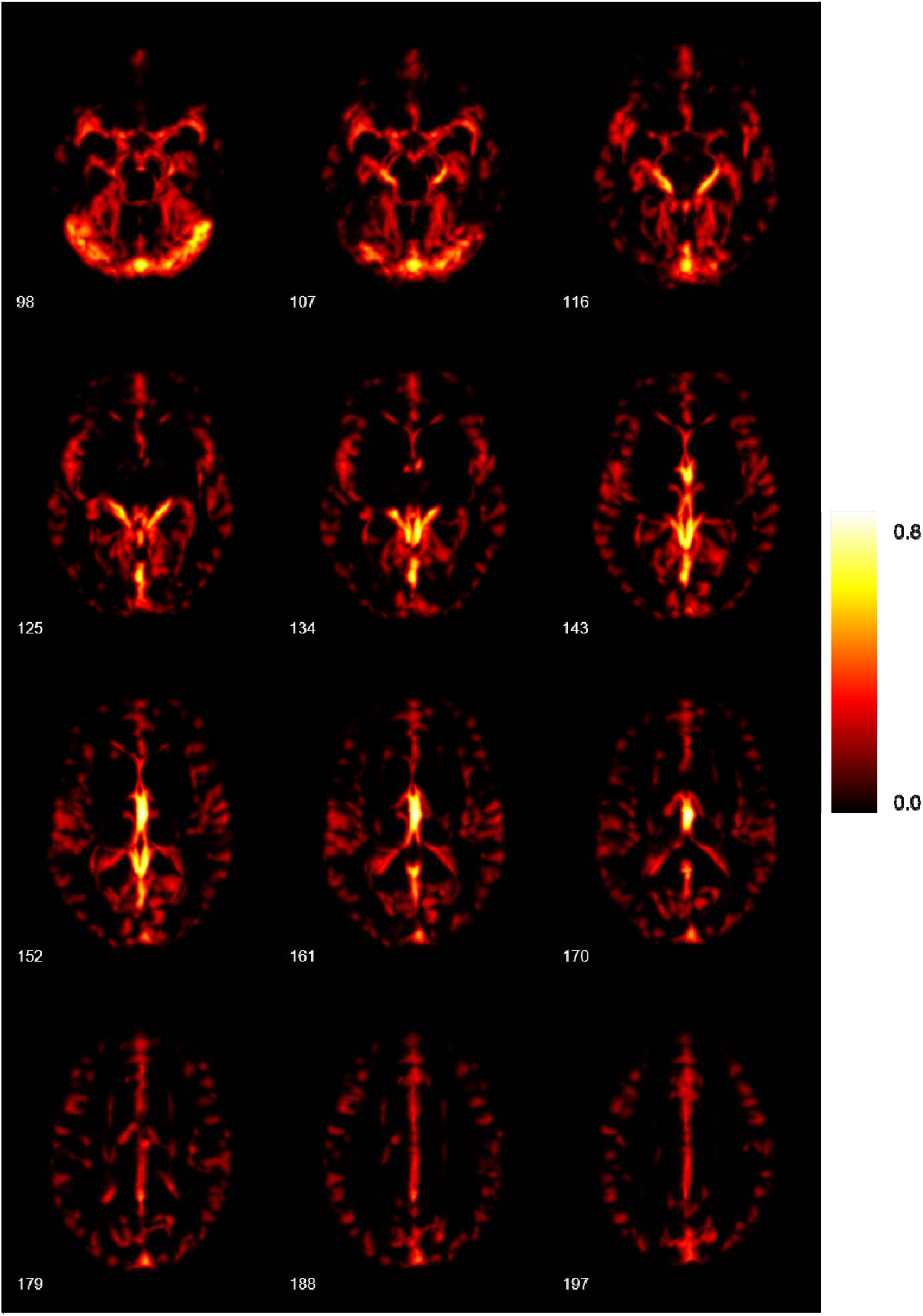
Maximum intensity projection across 20 slices. The images show the partial volume map of the multi-participant atlas from inferior to superior.

**Fig. 7:**
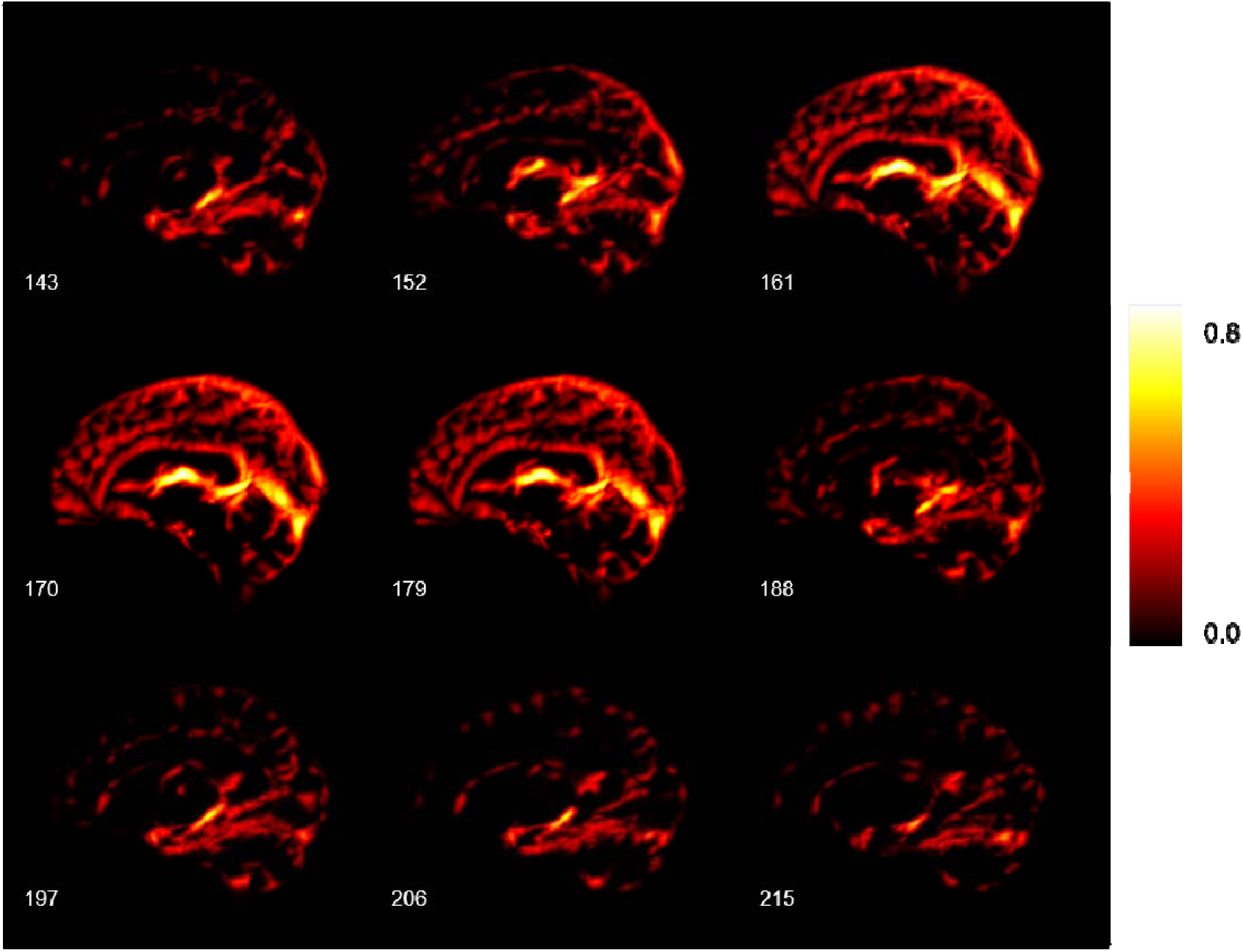
Maximum intensity projection across 20 slices. The images show the partial volume map of the multi-participant atlas from right to left.

**Fig. 8:**
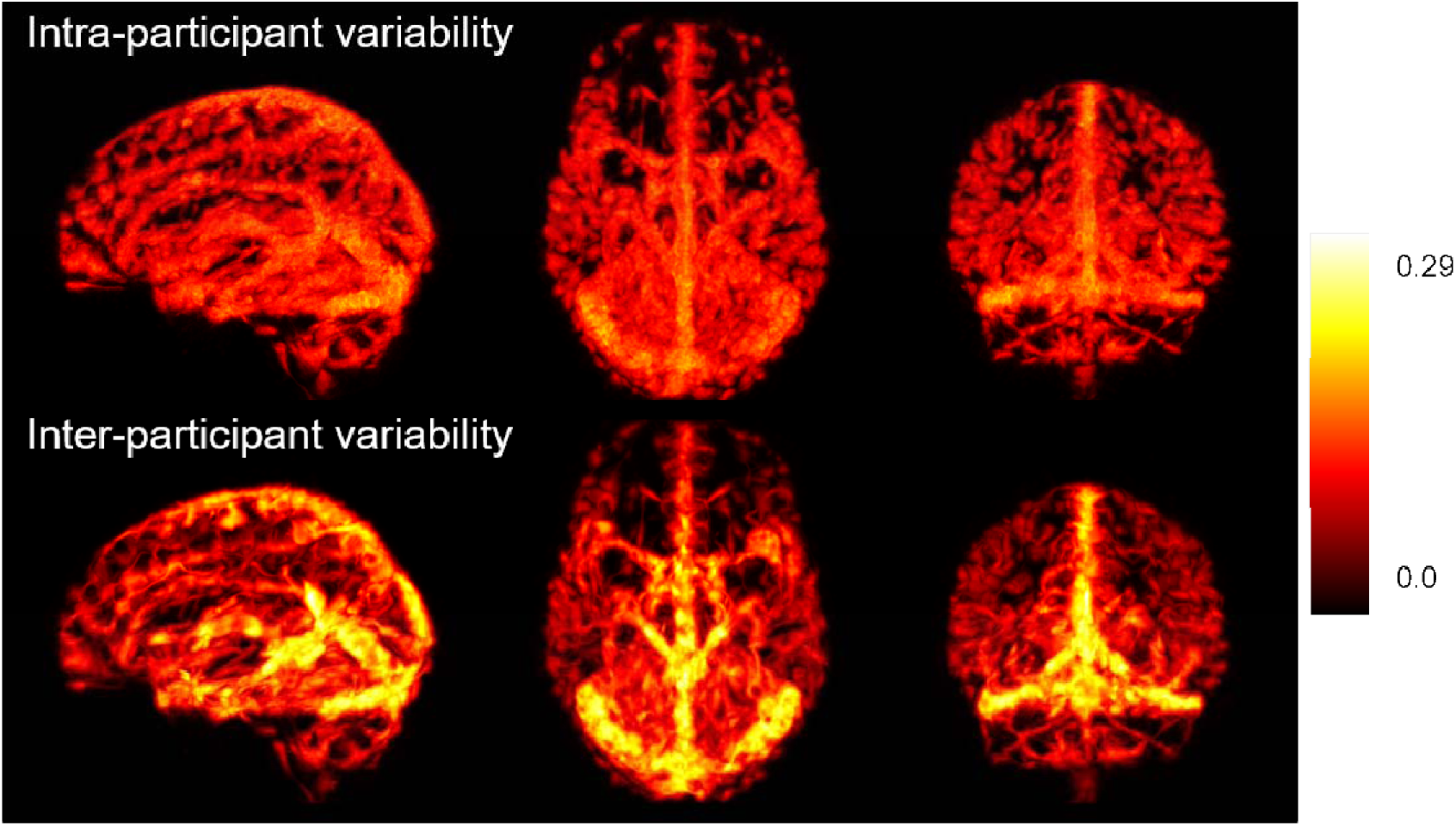
Maximum intensity projection over all slices; the images show the average standard deviation maps within participants (Intra-participant variability) and the standard deviation across participants (Inter-participant variability).

The voxel-based group probability density map is shown in **Fig. 9**, where 100% denotes that a voxel was classified as a vessel in all participants and time points. The regional densities of the MNI, the Harvard-Oxford and BASC064 atlases are shown in **Fig. 10**. Two density maps for the VENAT atlas were used, one with a higher sensitivity (voxels classified as veins in at least two time points) and with a higher specificity (voxels classified as veins in at least three time points). The region with the highest density across all three parcellation atlases was found to be the insula. Regional densities in the MNI atlas were found to be homogeneous (Occipital Lobe, Cerebellum, Parietal Lobe, Temporal Lobe and Frontal Lobe between 10.28 % ± 1.20 % to 11.94 % ± 1.07 %), with the highest density detected in the insula (15.09 % ± 1.06 %). The density distributions of the Harvard-Oxford atlas and the BASC064 were found to be more heterogeneous, however. The regional densities in the Harvard-Oxford atlas varied between 1.94 % ± 0.36 % / 2.91 % ± 0.69 % for the frontal pole for the high specificity and high sensitivity values respectively and 24.31 % ± 1.99 % / 29.51 % ± 2.30 % for Heschl’s Gyrus (including H1 and H2) for the high specificity and high sensitivity values respectively. Regional densities for the BASC064 atlas ranged between 0.20 % ± 0.26 % / 0.40 % ± 0.47 % for the temporal pole for the high specificity and high sensitivity values respectively and 17.18 % ± 2.45 % / 21.78 % ± 2.94 % for the lateral visual network anterior region for the high specificity and high sensitivity values respectively.

**Fig. 9:**
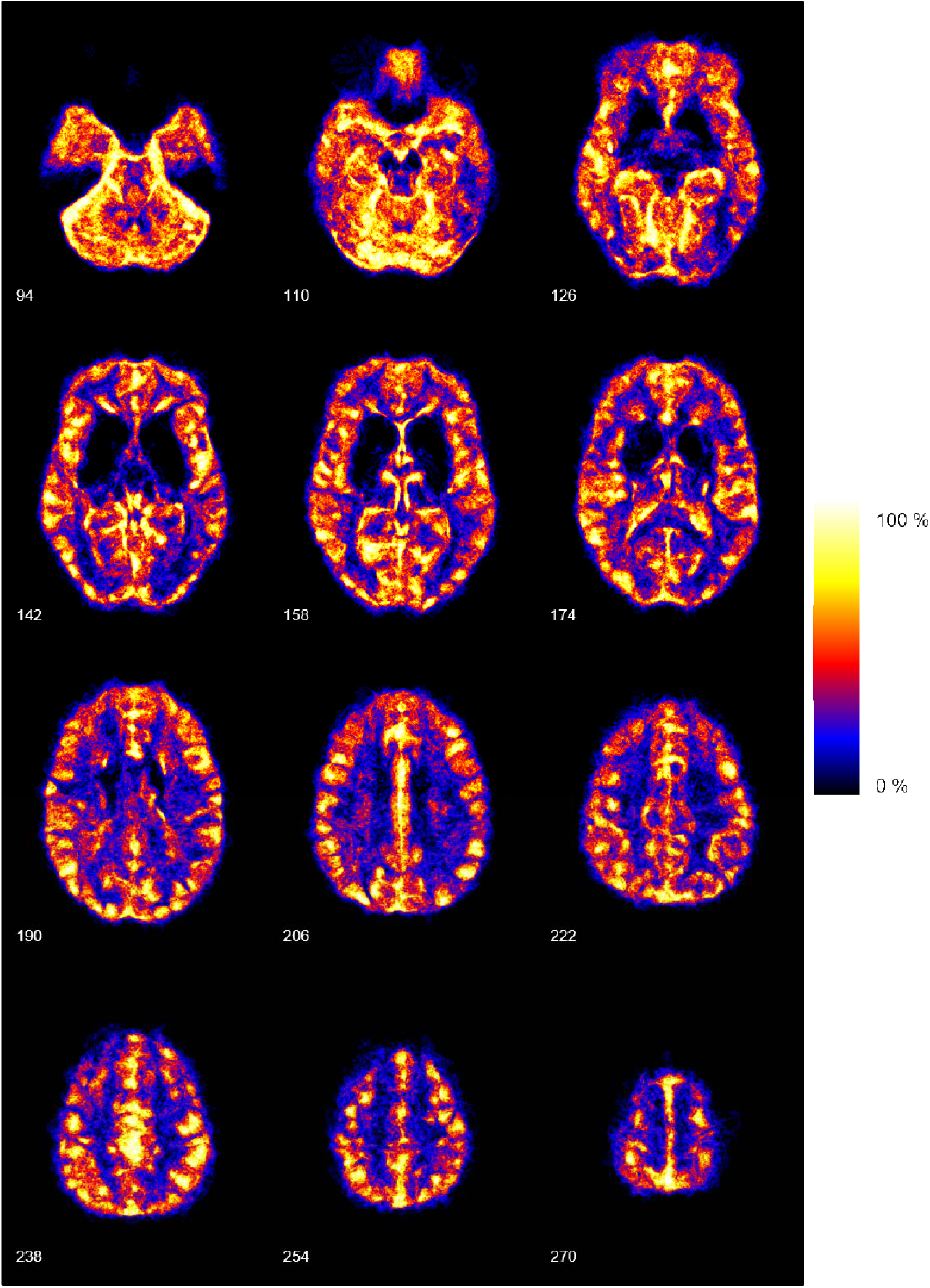
Individual axial slices of the probabilistic high sensitivity density atlas from inferior to superior.

**Fig. 10:**
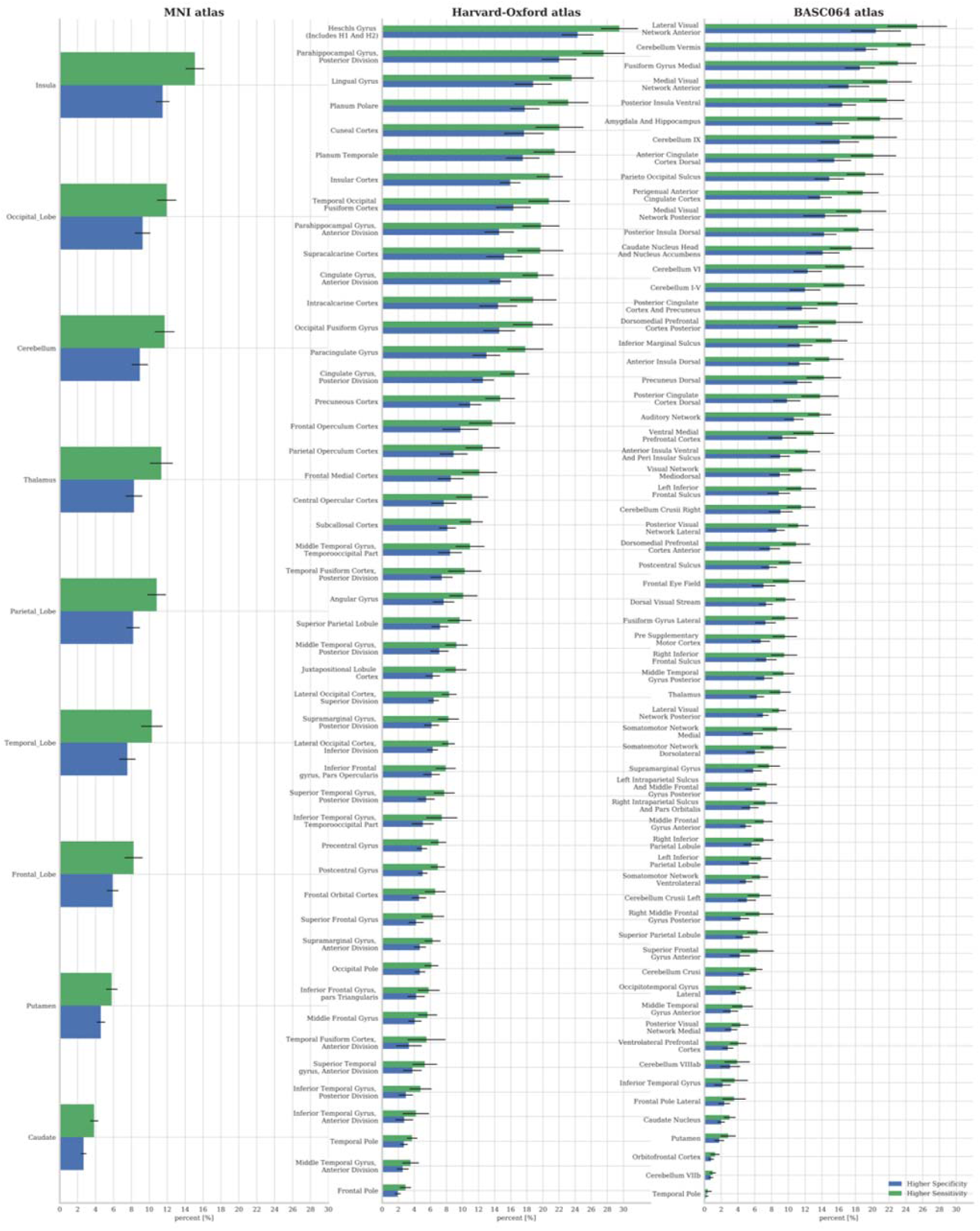
Regional density comparison between the higher sensitivity and higher specificity analyses in the MNI, Harvard-Oxford and BASC064 atlas.

Average diameter and curvature maps are shown in **Fig. 11**. Diameter and curvature maps from the multi-participant average were found to have a protracted range of values as compared to individual averages. However, the mean value of the average diameter map was similar to the individuals’ (mean: 0.84 ± 0.33 mm) but with increased standard deviation (mean: 0.26 ± 0.15 mm) reflecting inter-participant variability. Furthermore, the smallest diameter detected (0.4 mm) was larger than the theoretically detectable limit, as well as the values found in the individual averages. Average curvature of the VENAT atlas across the entire brain was similar to that of individual averages: 0.11 ± 0.05 mm^−1^. Standard deviation was also similar to the individual averages with an average value of 0.06 ± 0.03 mm^−1^.

**Fig. 11:**
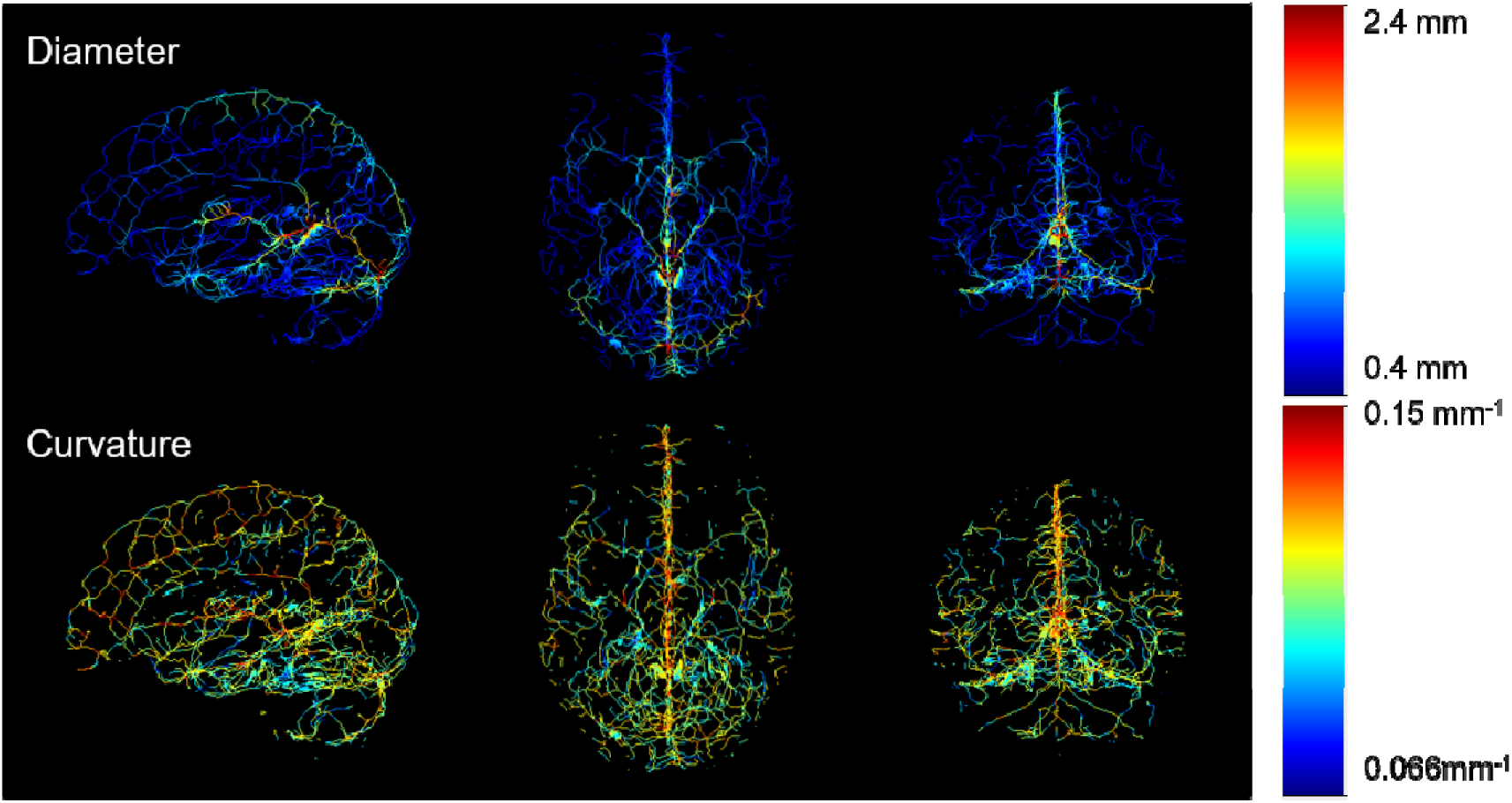
Average vessel diameter and curvature of the multi-participant atlas.

## Discussion

Here we present a high resolution atlas of the cerebral venous vasculature based on 7 T QSM in 20 young healthy individuals with five acquisitions each. This atlas was generated through multi-step registration of segmented vessels and therefore shows the average venous positions across participants. The inclusion of multiple QSM maps for each participant ensured a high-quality input vessel map for each participant for the subsequent across-participant registration. The atlas shows that vessels are consistent across time points within a single individual, leading to high quality individual vessel maps across days with low standard deviations. The multi-participant atlas highlights the high variability in vessel location, but especially in vessel diameter across individuals. The VENAT atlas also includes maps of the curvature of vessels, providing additional information about venous structure that has not previously been available. Overall, the VENAT atlas is the highest resolution whole-brain venous atlas currently available and will be an ideal tool for investigating the venous contribution of BOLD signals, as well as venous changes in aging and disease.

### Single participant average

To improve the quality of the individual inputs into the across-participant registration, we used five QSM acquisitions per participant. This was both to increase signal-to-noise ratio (SNR) and to reduce spurious vessels due to noise in our segmentations. Across-day individual averages for the LV and the HV participant, and the average standard deviation map of all participants illustrate the range of data quality used in this atlas. Participants with high SNR have a large number of systematically detected small vessels and relatively few regions where no vessels have been detected. Motion in some participants during the scan led to decreased venous contrast and SNR, resulting in a less consistent segmentation across days and more locally variable distribution of larger and blurrier vessels. This impact of data quality is also visible in larger diameter estimates and a larger standard deviation for noisier participants, highlighting the fact that the uncertainty added by lower SNR is expressed more strongly in the diameter estimates than in the position of vessels. This is likely due to the fact that we create a probability distance map of the skeleton to perform registration, so vessels that are not as well segmented across days were still registered into similar positions, but with uncertainty reflected in their exact diameter.

Curvature estimates appear largely uninfluenced by data quality differences, at least within this study which included only experienced 7 T participants. This can be seen from the identical curvature ranges and similar standard deviations for LV and HV. Therefore, curvature may be a valuable metric to use when comparing vessel structure across the lifespan or in disease populations, as data acquired in older adults and patients is likely to be noisier due to higher iron content (Wayne Martin et al. 1998; Zecca et al. 2004) and greater movement. Unfortunately, to our knowledge there is no ground truth quantification of the curvature expected from the human venous vasculature in vivo. While post-mortem work has shown curvature to increase with age and disease (Brown and Thore 2011), curvature is typically not quantified per se. Therefore, the validity of the estimates presented in this manuscript remains to be established. Nevertheless, the data presented here is a first step toward establishing normative values for QSM-based estimates of curvature.

### The VENAT vessel atlas

The individual averages across five days were then used to create the VENAT atlas from 20 participants, thus combining 100 measurements of the vasculature. The resulting atlas shows a prominent contribution of larger veins, indicating a lower variability in the position of larger veins. This is consistent with other atlasing work by Bernier et al. and Ward et al. (Ward et al. 2018; Bernier et al. 2018). Smaller veins, while abundant in the individual averages, are sparser in the across-participant average and the smallest diameter detected in the atlas (0.4 mm) is larger than the theoretical detection limit as well as the smaller vessels detected in the individual averages. This is due to the fact that smaller veins have to be consistent within a certain precision across all participants in order to be aligned. This sparsity of smaller veins in the overall atlas is present despite the higher field and spatial resolution used here, which allowed us to detect smaller cortical veins more consistently than in these previous atlases. This highlights one of the difficulties of venous atlasing, since the position and shape of smaller vessels is highly variable across individuals. However, this variability is still informative since if the VENAT atlas is taken as a normative atlas, then the standard deviation maps made available online can be used to detect abnormal venous structure due to disease.

The partial volume values in the atlas range from 0 to 0.8 (**Fig. 6** and **Fig. 7**), while PV in the individual participant averages include values between 0 and 1 (**Fig. 2**). This further emphasizes the variability across individuals, since this not only indicates that no voxel is entirely venous in the multi-participant average, but also that the variability across participants means that no vessel is ever completely consistent across participants. This is especially striking for larger veins which span more than one voxel, since even though registration of these vessels is somewhat straightforward, there is never any complete agreement on their exact location or diameter. Therefore, the atlas depicts the common location of veins and can be used to understand venous variability across participants. The multi-participant standard deviation maps demonstrate a high variability in vessel diameter especially for larger vessels, due both to the segmentation errors and inter-participant variability in vessel location that manifests as noise in the diameter estimate. The atlas also shows fewer vessels of small diameters as these are more variable and may not have been successfully registered across participants. This variability across participants in smaller veins especially is both consistent with other atlases and with existing post-mortem work (Duvernoy et al. 1981; Ward et al. 2018; Bernier et al. 2018).

In contrast to the homogeneous densities across the MNI atlas, the Harvard-Oxford atlas and the BASC064 atlas show heterogeneous densities (**Fig. 10**), probably due to the smaller size of the regions in these two atlases as compared to the MNI atlas. This heterogeneity of venous vessel density seen in the BASC064 and Harvard-Oxford atlas has also been described by Miyawaki et al., Vigneau-Roy et al. and Bernier et al. (Miyawaki et al. 1998; Vigneau-Roy et al. 2014; Bernier et al. 2018). The densities of the higher sensitivity analysis (vessels present in at least two time points) shown in **Fig. 10**, are almost identical to the vascular densities reported by Vigneau-Roy et al., whereas the lower density values of the higher specificity analysis (vessels present in at least three time points) are more similar to the values reported by Bernier et al. These similarities diverse densities may be due to the different thresholds used with the different multiscale vessel filter approaches of different studies. While Bernier et al. used a threshold of 0.5 to define their vessels, making it more similar to our more stringent analysis, Vigneau-Roy et al. used a very low threshold of 0.01, making it more similar to our higher sensitivity analysis. It is important to note however that the results for both the higher sensitivity and higher specificity analyses reported here were calculated without any threshold on the vessel filter. It is therefore likely that this apparent similarity in results does not stem from the same relationship between noise, resolution and consistency of vessel detection across days given the higher resolution used here and the greater number of repeated acquisitions. However, Bernier et al. reports a higher average density in the temporal gyrus, which may be due to signal drop-outs due to intra-voxel dephasing near air/tissue interfaces at higher field strength, i.e. 7 T (Olman and Yacoub 2011).

Regional analysis of vascular densities also revealed a pattern whereby primary sensory brain regions (visual area in the BASC064 atlas and Heschl’s Gyrus (auditory area) in the Harvard-Oxford atlas) followed by the insula (BASC064 and Harvard-Oxford atlas) showed some of the higher densities detected. This is consistent with data from post mortem and animal studies showing high vessel densities in these areas and reflecting their high degree of functional activity (Duvernoy et al. 1981; Bell and Ball 1985; Zheng et al. 1991).

The curvature atlas showed similar values and variability to the individual participant averages, largely due to the fact that these measures are extracted on the single participant level and averaged. Compared to vessel diameter, the measure appears more stable across participants. Validation with other techniques, including post-mortem work, would help determine whether the curvature values of the VENAT atlas correspond to true population averages or whether the measurements are impacted by vessel variability across individuals and represent underestimates of the true curvature. However, as no quantitative measurement of curvature in human cerebral veins could be found, no comparison was possible for this work.

### Other venous atlases

The recent increased interest in venous physiology has led to two other venous atlases being published recently (Ward et al. 2018; Bernier et al. 2018). The first atlas was presented by Ward et al. and is a probabilistic atlas based on combinations of QSM, SWI and manual venous segmentations. This atlas was created through averaging individual venous maps created from an optimized combination of these three types of images, but without explicitly registering segmented vessels. Rather, matrices from T1 to MNI152 registration were applied to these individual vascular trees. One advantage of this atlas is the fact that since all three types of images were used, the atlas includes the contribution of large veins which were obtained from SWI and the basal ganglia, which were obtained from manual segmentations. While our VENAT atlas necessarily excluded the deep gray matter, it is based on a larger group of participants, and a higher resolution image acquisition at higher field strength, allowing us to detect smaller vessels. Furthermore, by registering vascular trees rather than applying a T1 weighted-based registration on vessel trees, we are able to recover important information about venous structure such as average location, diameter and curvature. One difference between these two atlases is also the age range included. The atlas proposed by Ward et al. includes a wide age range (52.6 ± 25.2 years), while ours is based only on healthy younger adults (25.1 ± 2.5 years). While this may make the Ward et al. atlas more valid as a lifespan atlas, our new atlas may represent a more normative representation of the early, healthy vasculature, before any aging or disease-related changes have occurred. More studies on vascular changes in aging and disease are required to understand whether any changes can be observed in the range of diameters detectable using the VENAT atlas.

On the other hand, the probabilistic atlas by Bernier et al. differs both from the Ward et al. atlas and our atlas as it is both an arterial (based on time-of-flight angiography) and venous atlas (based on 3 T SWI imaging). Similarly, to the VENAT atlas, the atlas presented by Bernier et al. is based on non-linear registration of venous segmentations, also performed using ANTS. The age ranges used for the creation of both atlases are also similar. The VENAT atlas provides some advantages, including the higher field and resolution of the acquired data. We detected veins as small as 0.4 mm in diameter (Bazin et al. 2016) as opposed to the 0.6 mm limit of the atlas proposed by Bernier et al. Furthermore, the higher field strength provides a higher contrast, which further enhances our ability to detect smaller veins (Duyn et al. 2007). On the other hand, the use of SWI instead of QSM allowed Bernier et al. to map larger veins, since these appear with excellent contrast in SWI maps, but are shaved off at least partially in QSM when eroding the brain mask to avoid artifacts from air-tissue interfaces. However, the use of SWI may be suboptimal for detection of veins in the falx cerebri (Ward et al. 2018). Orientation effects on the SWI image (Fan et al. 2014) and inclusion of non-local sources in SWI may furthermore lead to overestimation of small vein diameters (Haacke et al. 2004; Ward et al. 2018).

Other advantages of the VENAT atlas include the multiple time points used to build the individual segmentations used as input into the atlas. This resulted in higher data quality for these individual inputs and lessened the need for denoising algorithms such as non-local means filter used by Bernier et al. This also ensured that the vessels included in the individual inputs were real vessels, reproducibly detected across days.

### Implications of our work

Venous atlases are a recent development, both for technical reasons and because the importance of venous physiology is increasingly recognized. One of the most obvious areas of application for these venous atlases is for calculating bias from larger veins in BOLD-based imaging. It has been estimated that the intravenous contribution to the BOLD signal at 3 T is 40 - 70 % (Boxerman et al. 1995; Donahue et al. 2011), but in addition to this, highly stringent thresholding of statistical maps may further enhance this effect by selecting only the most prominent, and therefore venous voxels. This is problematic both in terms of interpretation of the magnitude of the BOLD signal in response to a task, but also in localizing the underlying neuronal activity. Estimates based on this analysis by Turner of the Duvernoy atlas would indicate that the parenchyma drained by the smaller veins visible in our atlas (0.4 mm) can be up to 8 mm away from the location of the veins (Turner 2002). While there are techniques, such as spin echo BOLD fMRI, to minimize the intravascular venous contribution, they suffer from a lower sensitivity than gradient echo fMRI (Parkes et al. 2005) and are therefore less commonly used. An alternative approach would be to try to estimate this venous bias. The three atlases presented so far could all be used as priors in fMRI analysis to reduce or at least understand the impact of large veins on activation detection using gradient echo BOLD fMRI. The advantages of our atlas are related to its higher resolution, allowing the detection of bias from smaller draining veins more closely related to neural activity. Furthermore, venous diameter could be used to estimate the relative bias of differently sized veins, while curvature maps could help detect bias due to orientation as compared to the main magnetic field. Recent work by Gagnon et al. has suggested that orientation of the cortex in a given voxel may have an impact on the BOLD signal (Gagnon et al. 2015). This is likely to also apply to signal arising from draining veins and detected as areas of high activity in fMRI analyses.

The other application of this work is in helping us understand aging and disease-related changes in brain physiology. This could both be in understanding the changes in aging and disease-related biases in the BOLD signal, by identifying changes in venous structure that can impact BOLD signal location and amplitude biases due to draining veins, but also in comparing venous maps to our normative population. Aging, vascular diseases and dementia are associated with changes to the venous vasculature such as venous collagenosis (Brown et al. 2002) and CVT (Towbin 1973; Villringer et al. 1989; Einhäupl et al. 1991; Vogl et al. 1994). These could affect venous density, diameter and curvature and atlases such as the one proposed here could be used as normative data to attempt to detect and quantify these abnormal changes of density, diameter and curvature in vivo. The VENAT atlas cannot be generalized across age and disease, but can be used as normative data for clinical and research studies to better understand the influence of aging and diseases on the venous vasculature. Furthermore, the atlas can be used to estimate biases in BOLD measurements from large draining veins.

### Limitations

The main limitation of this work is related to resolution, as only larger veins (> 0.3 mm) can be measured. This is problematic as much of the bias in BOLD arises from smaller veins and much of the changes in aging and disease also occur in smaller veins (Brown and Thore 2011). This limited resolution is a consequence of limited acquisition time in live participants, as well as time limitations due to the impact of movement on QSM data quality. This limited resolution also makes validation of this work more difficult, as published post-mortem work such as the Duvernoy atlas (Duvernoy et al. 1981) only includes veins up to 380 µm. However, the higher field strength used for this atlas allowed the use of a higher resolution than that used for previous atlases. Future work using prospective motion correction and other advanced acquisition schemes may provide even higher resolutions for vascular atlasing.

Vessel segmentation based on QSM is also challenging for the largest veins and sinuses located at the surface of the brain, due to the masking required for the QSM reconstruction to avoid streaking artifacts from air-tissue interface. In this work, an erosion of 8 voxels was required to avoid these artifacts. As a result of this, the largest veins, such as the sagittal sinus, were not entirely included in the QSM images, leading to a greater variability in the detected diameter of the larger veins at the cortical surface across participants. In future versions of this atlas, we hope to remedy this problem by combining SWI images and QSM for the segmentation of the larger veins at the surface of the cortex, or by performing a more approximate QSM reconstruction for locating vessels at the surface of the cortex in spite of the artifacts (Topfer et al. 2015).

An additional aspect is the data quality and the reproducibility of the QSM reconstructions. Though the reproducibility of QSM maps at 3 T has been shown to be high (Deh et al. 2015), the reproducibility of vessel segmentations has never been assessed. Here we show that in high quality data, a consistent dense vessel tree can be reconstructed, but that even small variations in data quality can impair the ability to detect smaller veins or can lead to variations in the diameter estimation. Prospective motion correction of QSM acquisitions could lessen the impact of motion on data quality and render these techniques more robust. Even with highly reproducible vascular trees as in our best participants, the intrinsic variability of human vasculature is a challenge to atlasing. While we could show that careful alignment of the vascular tree across participants provides a detailed map for many veins, it is also clear that measures from individual segmentations are more informative than atlas-based trends, for instance when assessing biases in BOLD responses. However, when using existing or publicly available BOLD datasets that do not include SWI or QSM acquisitions, a normative atlas could still be instrumental for determining the impact of veins on population-level BOLD results.

### Conclusion

The VENAT atlas is the first atlas depicting veins using an ultra-high field MRI at 7 Tesla. It was built with a multi-step registration approach on segmented veins. An average image of each participant across multiple time points was used as an input for the atlas itself. The atlas consists out of four maps: partial volume, diameter, curvature, and venous density maps which are all freely available (https://figshare.com/articles/VENAT_Probability_map_nii_gz/7205960). The VENAT atlas is a promising tool that can be used as normative data for clinical and research studies to better understand the influence of aging and diseases on the venous vasculature, but also to estimate biases in BOLD measurements from large draining veins.

## Acknowledgement

The authors thank Domenica Wilfling and Elisabeth Wladimirov for their help with data acquisition and logistics of the multi-modal plasticity initiative (mMPI) dataset. This work was supported by the Max Planck Society, the Canadian National Sciences and Engineering Research Council (RGPIN-2015-04665, C.J.G.), the Heart and Stroke Foundation of Canada (N.I.A. C.J.G.), the National Institute of Health (1K99NS102884, A.P.F.) and the Quebec Bio-Imaging Network (QBIN) for the scholarship for Training course abroad (J.H.).

